# Role of the pre-initiation complex in Mediator recruitment and dynamics

**DOI:** 10.1101/207282

**Authors:** Elisabeth R. Knoll, Z. Iris Zhu, David Landsman, Randall H. Morse

**Affiliations:** Department of Biomedical Sciences, University at Albany School of Public Health, Albany, New York, 12201-0509; Wadsworth Center, New York State Department of Health, Albany, New York 12201-0509; Computational Biology Branch, National Center for Biotechnology Information, National Library of Medicine, NIH, Bethesda, MD 20814

**Author notes:** Correspondence: Randall H. Morse.

**Keywords:** Transcription, yeast, Mediator, TATA-binding protein, Tafs

## Abstract

The Mediator complex functions in eukaryotic transcription by stimulating the cooperative assembly of a pre-initiation complex (PIC) and recruitment of RNA Polymerase II (Pol II) for gene activation. The core Mediator complex is organized into head, middle, and tail modules, and in budding yeast (*Saccharomyces cerevisiae*), Mediator recruitment has generally been ascribed to sequence-specific activators engaging the tail module triad of Med2-Med3-Med15 at upstream activating sequences (UASs). We show that *med2*Δ *med3*Δ *med15*Δ yeast are viable and that Mediator lacking Med2-Med3-Med15 is associated with active promoters genome-wide. To test whether Mediator might alternatively be recruited via interactions with the PIC, we examined Mediator association genome-wide after depleting PIC components. We found that depletion of Taf1, Rpb3, and TBP profoundly affected Mediator association at active gene promoters, with TBP being critical for transit of Mediator from UAS to promoter, while Pol II and Taf1 stabilize Mediator association at proximal promoters.

## Introduction

The Mediator complex plays a central, highly conserved role in eukaryotic transcription by stimulating the cooperative assembly of a pre-initiation complex (PIC) and recruitment of RNA Polymerase II (Pol II) for gene activation (1–9). In budding yeast (*Saccharomyces cerevisiae*), Mediator is composed of 25 subunits divided into four domains based on structural and functional criteria: the tail, middle, head, and kinase modules (10–12). Mediator recruitment has generally been ascribed to sequence-specific activators engaging the tail module triad of Med2-Med3-Med15 (13–15); consistent with this model, the tail module can under some circumstances be recruited independently of the remainder of Mediator (1, 15–19). Normally, however, the tail is recruited as a complex with the head, middle, and kinase modules at upstream activation sequences (UASs) (8, 20–22). Once Mediator is recruited, the kinase module dissociates from the complex and Mediator bridges the enhancer to connect with Pol II and general transcription factors (GTFs) at the proximal promoter of active genes (4, 18). Mediator association with the PIC at proximal promoters is quickly disrupted following phosphorylation of the Pol II C-terminal domain (CTD) by Kin28, marking the end of initiation (20, 21).

This orchestrated recruitment of Mediator is broadly required for transcription, evidenced by studies that showed a loss of Mediator function results in greatly decreased mRNA levels and concomitant diminished association of Pol II with essentially all Pol II transcribed genes (1, 17, 20, 23–25). In spite of the purported role of the tail module triad in Mediator recruitment by transcriptional activators, *med3Δ med15Δ* yeast are viable, show transcriptional defects for only 5-10% of active genes during normal growth, and exhibit continued Mediator and Pol II occupancy by ChIP-seq at most proximal promoters (4, 17, 18, 26). Additional studies report that the tail module is not required for Mediator’s interaction with the PIC (4, 18, 24). That transcription is unaffected at many genes in *med3Δ med15Δ* yeast supports the notion that Mediator is recruited through an unknown mechanism that is independent of the tail module, possibly through components of the PIC which are known to engage Mediator subunits at proximal promoters (2, 3, 27–29). Indirect recruitment of Mediator via interactions with the general transcription machinery has also been proposed on the basis of *in vitro* experiments (30).

Several studies show that a loss of Mediator leads to a decrease in PIC components and diminished Pol II presence at active genes, highlighting the unilateral importance of Mediator in stimulating the assembly of a PIC (16, 17, 23, 25, 31–33). Yet, how the PIC itself influences Mediator recruitment has received little attention (28, 34). Mediator’s interaction with the PIC at proximal promoters is likely brief in light of recent evidence demonstrating that Mediator association at proximal promoters is transient, making interactions with the PIC difficult to assess (20, 21, 35). The close spatial relationship of Mediator with the PIC at proximal promoters has been shown in single particle cryo-electron microscopy and cross-linking mass spectrometry studies of yeast in which Mediator is bound with the PIC (24, 34, 36, 37). The TFIID-TBP complex and Pol II in particular occupy proximal promoters and have the potential to engage Mediator directly.

To investigate PIC-dependent Mediator recruitment *in vivo* we have performed ChIP-seq against Mediator subunits in *Saccharomyces cerevisiae* after employing the anchor away method of nuclear depletion to export individual components of the PIC to the cytoplasm (38). We confirm the reported effect of Mediator stabilization at proximal promoters in the *kin28*-AA strain. In the same background, we rigorously show continued tail-less Mediator association at proximal promoters after complete loss of the tail module triad in *med2Δ med3Δ med15Δ* yeast. We also compare Mediator association in *tbp*-AA and *rpb3*-AA strains by ChIP-seq to assess the individual contribution of TBP and Pol II on Mediator occupancy. Mediator occupancy at proximal promoters was also evaluated after the double depletion of Kin28 with Taf1, TBP, or Rpb3. Taken together, our findings provide new insights into mechanisms of Mediator recruitment during gene activation in yeast.

## Results

### Mediator is recruited to gene promoters in the absence of the tail module triad

The Mediator tail module triad, comprising Med2, Med3 (Pgd1), and Med15 (Gal11) (39), has been implicated as a primary target of activators in yeast (2, 14, 15). However, previous work has shown that yeast lacking any two of the three tail module triad subunits are viable (15), and gene expression and Mediator recruitment are relatively unaffected for most genes in *med3*Δ *med15*Δ yeast (4, 18, 26). Med2 recruitment is substantially impaired in *med3*Δ *med15*Δ yeast (4, 17, 18, 26), so the modest effects on mRNA levels in these yeast raise questions about the role of the tail module triad in Mediator function genome-wide. However, Med2 recruitment is not completely abolished in *med3*Δ *med15*Δ yeast (Fig. 1A), and Mediator association with promoters is only modestly diminished in *med3*Δ *med15*Δ mutant strains (4, 18). Therefore, to assess more rigorously the role of the tail module triad in Mediator function, we sought to determine the effect of deletion of all three tail module triad subunits on yeast viability and Mediator recruitment.

**Figure 1.**
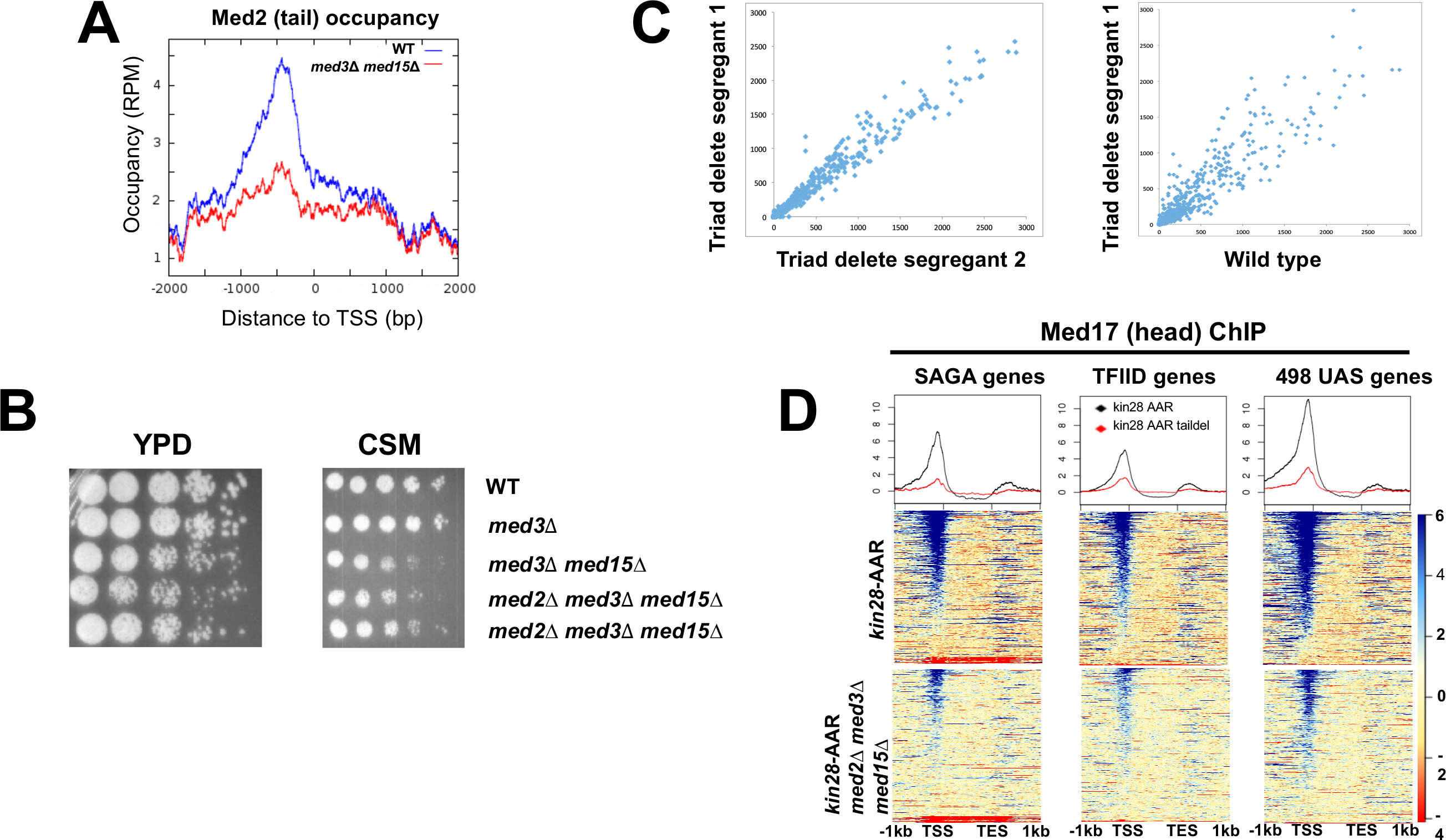
Mediator recruitment in *med2*Δ *med3*Δ *med15*Δ yeast. (A) Med2 occupancy, in reads per million, in wild type and *med3*Δ *med15*Δ yeast, averaged over 498 genes that exhibit Mediator signal at UAS regions in wild type yeast (4). Data from Paul et al. (17). (B) Spot dilutions (five-fold from left to right) of the indicated strains were made on YPD or CSM plates and allowed to grow at 30°C for 3 days. (C) Comparison of transcript levels (reads per million) by RNA-seq for two independent *med2*Δ *med3*Δ *med15*Δ segregants (left panel), and between segregant 1 and the wild type parent strain (right panel). Only data for mRNA genes are plotted, after normalization to total mRNA. (D) ChIP-seq was performed against Med17 in *kin28-AA* and *med2*Δ *med3*Δ *med15*Δ *kin28-AA* yeast after 1 hr treatment with rapamycin (designated “AAR”). Reads were mapped to all SAGA-regulated genes, all TFIID-regulated genes, and to the 498 genes exhibiting detectable Mediator ChIP signal at UAS regions in wild type yeast (4). Genes were normalized for length and aligned by transcription start site (TSS) and transcription end site (TES).

Segregants from yeast heterozygous for *MED2*, *MED3*, and *MED15* (complete ORF deletions) included viable *med2*Δ *med3*Δ *med15*Δ mutants, which showed similar growth defects to *med3*Δ *med15*Δ yeast (Fig. 1B). RNA-seq showed that two independent segregants exhibited genome-wide expression patterns much more similar to each other than to wild type yeast (Fig. 1C). Down-regulated genes were enriched for SAGA-regulated promoters (75 of 164 genes down-regulated by > 2X in both segregants; p < 10^−34^, hypergeometric test), while up-regulated genes showed marginal enrichment (35 of 442 genes, p = 0.04), consistent with previous findings (26). RNA-seq also confirmed the triple deletion phenotype, with no reads present for sequences from the *MED2, MED3*, or *MED15* ORF.

We next examined the effect of loss of the tail module triad on Mediator recruitment. Mediator association with UAS elements is seen in ChIP experiments at only a fraction of active yeast genes, and its association with active promoters appears to be transient, making it difficult to assess Mediator association with gene regulatory regions in wild type yeast (17, 20, 21, 35, 40). Mediator ChIP signal at promoters is greatly enhanced under conditions in which phosphorylation by the Pol II CTD by the Kin28 subunit of TFIIH is prevented, apparently by inhibiting escape of Pol II from the promoter proximal region (20, 21). We therefore constructed a Kin28 anchor away (*kin28-AA*) strain harboring the *med2*Δ *med3*Δ *med15*Δ mutations to assess Mediator occupancy genome-wide. Treatment of this strain with rapamycin tethers Kin28 to the Rpl13A ribosomal subunit via FKBP12 and FRB tags on the respective proteins, and leads to Kin28 eviction from the nucleus during ribosomal protein processing (38). The strain also includes the *tor1-1* and*fpr1*Δ mutations, which confer resistance to rapamycin (38).

We used ChIP-seq to compare occupancy by Med17 (Srb4) (39) from the Mediator head module at SAGA-dependent and TFIID-dependent promoters (41). Consistent with previous work, treatment of *kin28-AA* yeast harboring intact Mediator allowed detection of myc-tagged Med17 at the majority of promoters (Fig. 1D) (20, 21), while a parallel experiment using an untagged strain, or an input sample, gave rise to very low signal (Fig. S1A). Mediator peaks observed after depletion of Kin28 corresponded well with those identified in a previous study (Fig. S1B) (20). In *kin28-AA* yeast lacking the Mediator tail module triad, Med17 ChIP signal was reduced but still detectable at the same promoters (Fig. 1D). The reduction in Med17 occupancy was proportionally larger at SAGA-regulated genes than at TFIID-regulated genes, possibly reflecting the enrichment of SAGA-regulated genes among genes depending most strongly on the tail module triad for transcription (Fig. 1D) (26).

Examination of a set of 498 genes for which Mediator association was detected at UAS regions in *KIN28*-WT yeast (4) indicates Med17 occupancy of promoters and UAS regions in *kin28-AA* yeast having an intact tail module triad (Fig. 1D, upper right panel). Mediator head module subunits exhibit weaker ChIP signal at many UAS regions than do tail module subunits (4, 18); nonetheless, Mediator occupancy was evident at many UAS regions, as seen in the heat map and the shoulder on the upstream side of the line graph in Fig. 1D, upper right panel. This occupancy appears to be reduced to baseline levels in *med2*Δ *med3*Δ *med15*Δ yeast, as was reported previously for *med3*Δ *med15*Δ yeast (4, 18).

Based on the effects on viability, genome-wide expression, and Mediator recruitment in *med2*Δ *med3*Δ *med15*Δ yeast shown here, we conclude that the tail module triad is essential for Mediator recruitment to UAS regions but is dispensable for most gene expression and for Mediator recruitment (albeit at reduced efficiency) to promoters of the large majority of active genes.

### Mediator association at UAS regions is stabilized by loss of TBP or Pol II

Results from Figure 1 and previous work indicate that while the tail module triad of Mediator is required for its association with UASs, Mediator can associate with many promoters independently of the tail module triad (4, 18). Based on known interactions of Mediator head and middle subunits with the general transcription machinery (2, 3, 11, 24, 27, 28, 34, 36, 37), we had previously suggested that Mediator might be recruited to tail module-independent targets via interactions with PIC components (26, 42). To examine this possibility, we monitored Mediator subunit occupancy genome-wide before and after depletion of PIC components using the anchor away method (38).

We chose two PIC components to subject to depletion: TATA-binding protein (TBP), and Rpb3, an essential subunit of Pol II. Depletion of either TBP or Rpb3 resulted in substantial loss of Pol II association, monitored by ChIP-seq using an antibody against Rpb1, as expected (Fig. S2). We examined Mediator binding before and after depletion of TBP and Rpb3 using Myc-tagged Med15, from the tail module triad, and Med18, from the head module. We first examined a set of 498 genes previously identified as displaying Mediator peaks at UAS regions (4), as Mediator ChIP signal at promoters is generally low due to rapid turnover during transcription (20, 21). In agreement with prior reports, we observed association of both Med15 and Med18 at UAS regions for many of these genes, while a shift to the promoter region was seen for both Mediator subunits in *kin28-AA* yeast after rapamycin treatment (Fig. 2A).

**Figure 2.**
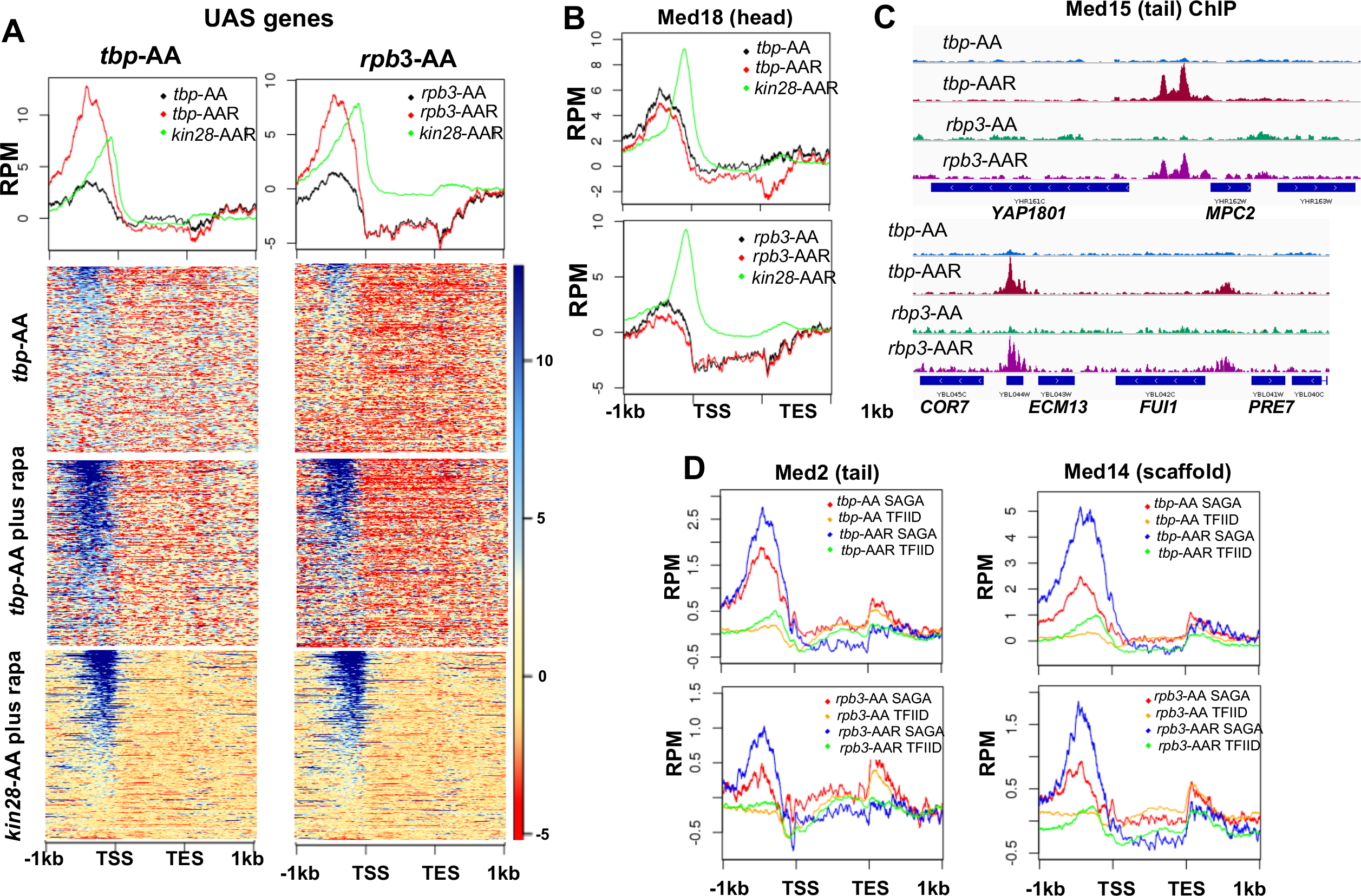
Effect of depletion of TBP and Rpb3 on Mediator occupancy. ChIP-seq was performed against (A) Med15 and (B) Med18 in *tbp*-AA and *rpb3*-AA yeast with (“AAR”) and without (“AA”) rapamycin treatment, and in *kin28*-AA yeast after rapamycin treatment. Heat maps (for Med15 ChIP) and line graphs (normalized reads per million, RPM) summarize results seen at 498 genes exhibiting detectable Mediator ChIP signal at UAS regions in wild type yeast (4). (C) Browser scans showing occupancy of Med15 at two chromosomal regions in *tbp*-AA and *rpb3*-AA yeast with and without rapamycin treatment. *YBL044W*, between *COR7* and *ECM13*, is an uncharacterized ORF and likely contains a UAS, as has been observed at many other uncharacterized or dubious ORFs (17). (D) Med2 and Med14 occupancy measured by ChIP-seq with and without depletion of TBP or Rpb3, at SAGA-regulated and TFIID-regulated genes.

Depletion of TBP resulted in a marked increase in Med15 association with UAS regions, while little effect was seen on the association of Med18 (Fig. 2A-C, and Fig. S2). Similar results were seen upon depletion of Rpb3 (Fig. 2A-C, and Fig. S2). ChIP-seq against another tail module triad subunit, Med2, yielded results similar to those seen with Med15, showing increased association after depletion of TBP or Rpb3, as did the scaffold subunit Med14 (Fig. 2D).

The increased ChIP signal for Med15 observed at UAS regions upon depletion of TBP or Rpb3 suggested that PIC component depletion might result in Mediator peaks being observed at UAS regions where they are normally difficult to detect. To test this idea, we examined Med15 ChIP-seq signal over all genes having transcription level > 32 (43), after removing the 498 genes identified as displaying Mediator UAS peaks (about 1400 genes) (Fig. 3A). Indeed, this analysis revealed several hundred genes that showed increased Mediator occupancy in their UAS regions upon TBP depletion where negligible occupancy was observed by ourselves or others under normal growth conditions (4). Gene ontology (GO) analysis of the 200 genes from this group showing the greatest increase in Med15 occupancy revealed a strong enrichment for ribosomal protein (RP) genes, and a direct examination of RP genes showed increased Mediator occupancy at UAS regions upon depletion of TBP (Fig. 3B).

**Figure 3.**
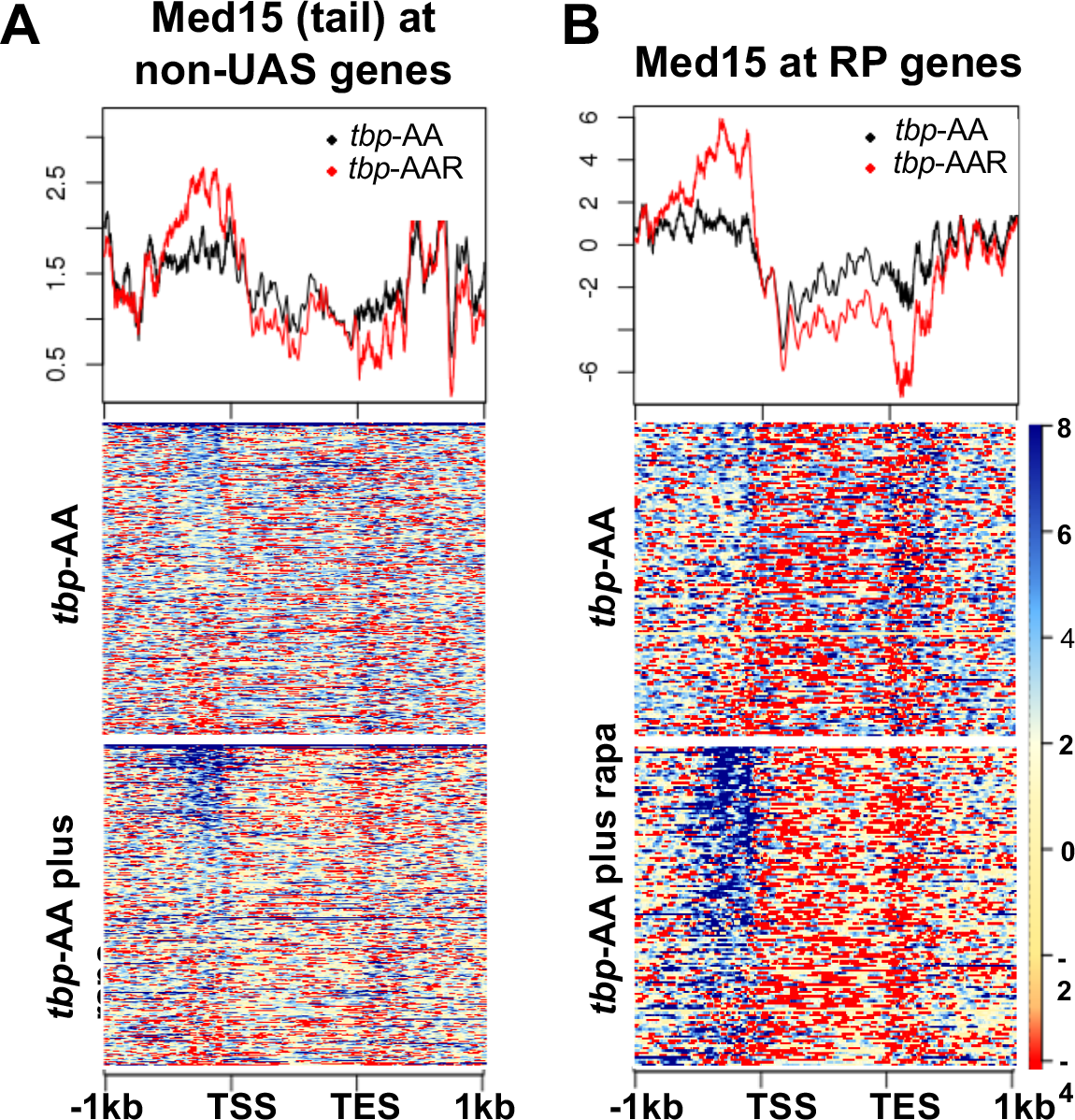
Effect of depletion of Taf1 on Mediator occupancy. (A) Med15 occupancy measured by ChIP-seq at the 2000 most highly transcribed genes, after removal of the 498 genes exhibiting detectable Mediator ChIP signal at UAS regions in wild type yeast (4), in *tbp*-AA yeast with and without rapamycin treatment. (B) Med15 occupancy measured by ChIP-seq at ribosomal protein (RP) genes in *tbp*-AA yeast with and without rapamycin treatment.

Our initial expectation was that Mediator occupancy would be *decreased* upon PIC depletion. Instead, we observed a clear increase in Med15 occupancy of UAS regions upon depletion of TBP or Rpb3. Furthermore, we previously reported decreased Med18 occupancy upon inactivation of Pol II using an *rpb1-1 ts* mutant (44). In the latter experiments, however, Med18 occupancy was measured at core promoter regions, not at UAS regions, and the observed decrease in ChIP signals at the restrictive temperature may have been misleading, due to the possibility of artifactual ChIP signals at actively transcribed open reading frames (45, 46).

A possible explanation for increased Med15 occupancy at UAS regions upon depletion of TBP or Rpb3 is that interruption of the transcription cycle by PIC disruption prevents the transit of Mediator from UAS regions to promoters and off the DNA. Inactivation of Kin28 prevents escape of Pol II, resulting in Mediator stabilization at gene promoters (20, 21). At genes for which Mediator association with UAS regions is observed, a single Mediator complex bridges the UAS and the promoter, with tail module subunits contacting the UAS and head and middle subunits engaging the promoter region through their contacts with Pol II and GTFs (4, 18). Interruption of the latter contacts could strand Mediator at the UAS. The increase in ChIP signal observed for Med15, but not for Med18, would be consistent with the most immediate contacts with UAS-bound activators occurring through tail module subunits (such as Med15), while ChIP signal for Med18 at UAS regions would depend on protein-protein cross-links, thus lowering the efficiency of ChIP and possibly making increased occupancy more difficult to detect. Alternatively, it is possible that the tail module, which can associate with UAS regions independently of the rest of Mediator under some conditions (1, 15–17), may bind to UAS regions independently of the head module at least some of the time (19).

Although this explanation is consistent with recent models of Mediator function at UAS regions (4, 18), the foregoing results do not help clarify how Mediator is recruited to promoters at the many genes for which Mediator association is not readily observed and for which expression does not depend on the Mediator tail module triad. We therefore next modified our approach to allow PIC component depletion under conditions in which Mediator association with active gene promoters was stabilized.

### Mediator association with promoters depends on PIC components

We next asked how PIC component depletion affected Mediator association under conditions in which Mediator binding to proximal gene promoters was stabilized. For this purpose, we constructed double anchor away strains in which both Kin28 and either TBP, Taf1, or Rpb3 were tagged with FRB to allow nuclear depletion upon treatment with rapamycin. We then separately introduced Myc-tagged Med15 and Med18 into each double anchor away strain, and performed ChIP-seq to examine occupancy of Med15, Med18, and Pol II after one hr of rapamycin treatment.

We first examined the effect of depletion of PIC components on Mediator occupancy at all genes, separated into SAGA-regulated and TFIID-regulated genes (41). Our rationale was that genes that depend on the tail module triad for transcription are enriched for SAGA-regulated genes, relative to TFIID-regulated genes (26); therefore, since most TFIID-regulated genes are transcribed independently of the Mediator tail module triad, they might show greater dependence on PIC components (42).

Depletion of either Taf1 or Rpb3 resulted in decreased occupancy of Med15 and Med18 in comparison to depletion of Kin28 alone at almost all TFIID-regulated genes and at the majority of SAGA-regulated genes (Fig. 4A-C). The effect of Taf1 depletion on Mediator occupancy at SAGA-regulated genes was unanticipated, but is consistent with Taf1 associating with and functioning in the transcription of SAGA-regulated as well as TFIID-regulated genes (47, 48). The greater occupancy seen after depletion of Taf1 or Rpb3 at SAGA-regulated genes compared to TFIID-regulated genes may be partly attributed to the stronger ChIP signal seen at a small fraction of SAGA-regulated genes (compare uppermost parts of heat maps for *kin28-AA* samples between SAGA-regulated and TFIID-regulated genes). However, a comparatively greater loss of ChIP signal is also seen when examining genes having similar levels of transcription (Fig. S3), and examples can also be found of individual TFIID-regulated genes exhibiting greater loss of Mediator ChIP signal than at SAGA-regulated genes (Fig. 5 and S4). We conclude that under conditions of Kin28 depletion, Mediator occupancy at gene promoters, particularly at TFIID-regulated genes, depends strongly on both Taf1 and Pol II.

**Figure 4.**
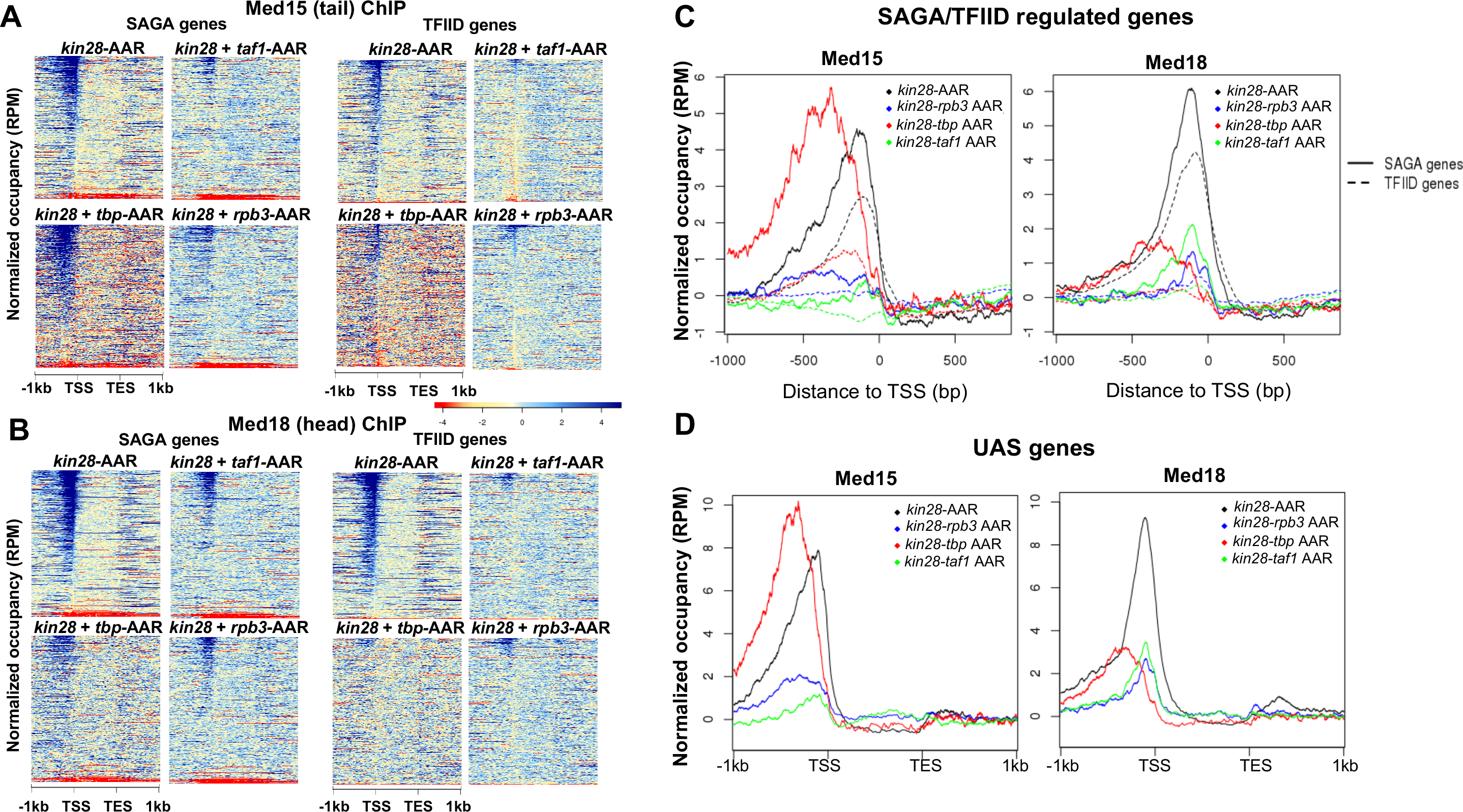
Effect of depletion of PIC components together with Kin28 on Mediator occupancy. (A) Heat maps showing occupancy of Med15 after depletion of Kin28 alone, or in combination with Taf1, TBP, or Rpb3, at SAGA-regulated and TFIID-regulated genes. (B) Same as (A), for Med18 occupancy. (C) Line graphs (normalized reads per million) depicting Med15 and Med18 occupancy after depletion as in (A) and (B), aligned at the TSS and shown for SAGA-regulated (solid lines) and TFIID-regulated (dashed lines) genes. (D) As in (C), but for genes exhibiting detectable Mediator ChIP signal at UAS regions in wild type yeast (4).

**Figure 5.**
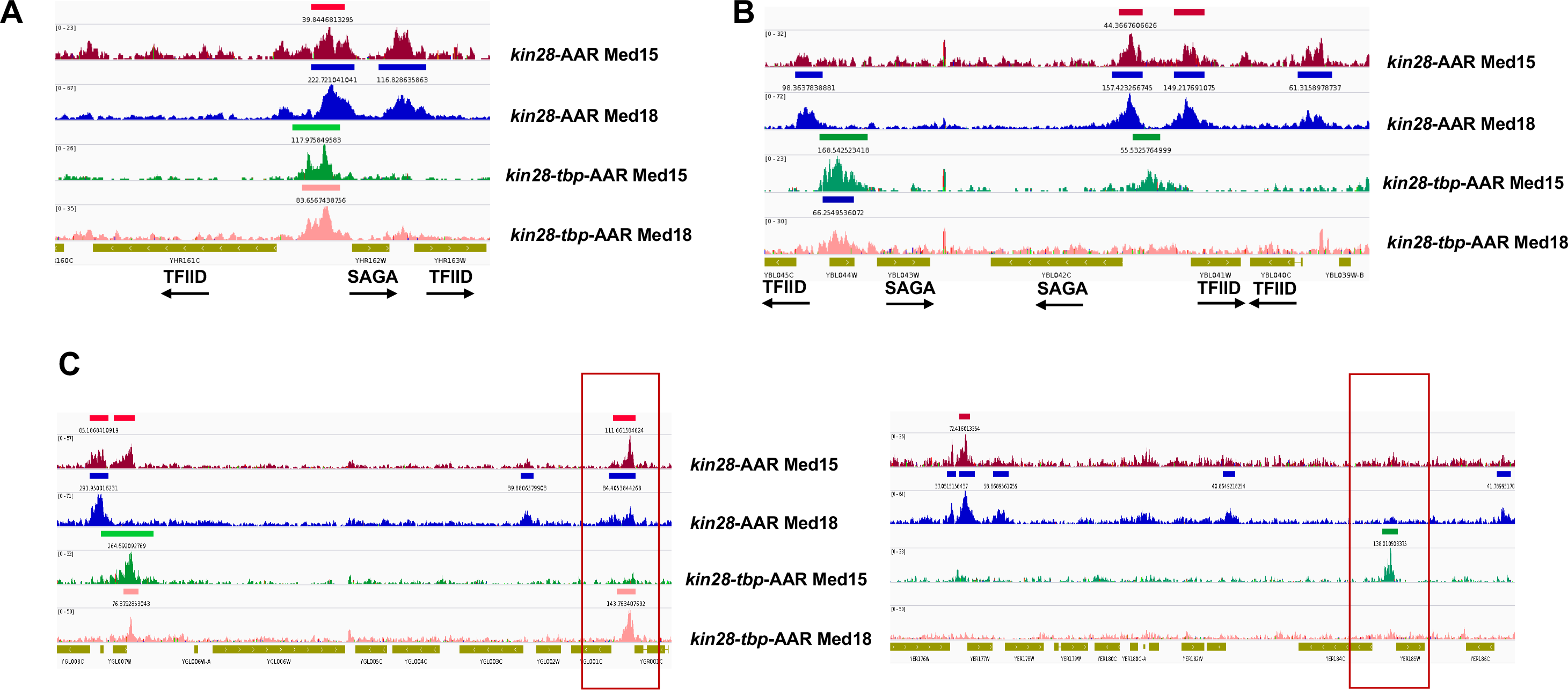
TBP depletion shifts Mediator occupancy from promoter to UAS sites. Browser scans showing Med15 and Med18 occupancy in *kin28*-AA and *kin28-tbp*-AA yeast treated with rapamycin, as indicated. (A) Upstream shift of Mediator at a SAGA-regulated gene and loss of occupancy at a TFIID-regulated gene upon depletion of TBP. Arrows depict direction of transcription. (B) Loss of Mediator peaks at promoters and gain of Mediator peaks at upstream regions at both TFIID and SAGA-regulated genes. *YBL044W* is an uncharacterized ORF and likely contains a UAS, as has been observed at many other uncharacterized or dubious ORFs (17). (C) At some loci, new Mediator peaks are visible after dual depletion of Kin28 and TBP (boxed regions).

### Depletion of TBP, and to lesser extent Pol II, stabilizes Mediator occupancy at UAS regions

The effect of depleting TBP together with Kin28 was markedly different from that of depleting Taf1 and Kin28 (Fig. 4A-C). At SAGA-regulated genes, occupancy of Med15 from the Mediator tail module was not decreased, but rather shifted upstream in *kin28-TBP-AA* yeast compared to *kin28-AA* yeast (Figure 4A & C). At TFIID-regulated genes, depletion of TBP also resulted in an upstream shift, along with a reduction in the occupancy of Med15 (Fig. 4A & C). Occupancy of Med18, from the head module, also shows a shift upstream at SAGA-regulated genes upon TBP depletion, but in this case the shift is accompanied by a reduction in signal, while Med18 occupancy is nearly eliminated at TFIID-regulated genes upon TBP depletion (Fig. 4B-C).

Depletion of Rpb3 together with Kin28 affected Mediator occupancy in a manner intermediate between effects seen in *kin28-TBP-AA* and *kin28-taf1-AA* yeast. As with depletion of Taf1, depletion of Rpb3 strongly decreased occupancy of both Med15 and Med18, especially at TFIID-regulated genes (Fig. 4A-C). However, in contrast to effects seen in *kin28-taf1-AA* yeast and similar to the effect of depleting TBP and Kin28, a shift in Med15 occupancy to more upstream regions (together with an overall decrease in occupancy) was seen at SAGA-regulated genes upon depletion of Rpb3 and Kin28 (Fig. 4A & C). This intermediate phenotype is also apparent upon analysis of the 498 genes at which Mediator occupancy is observed in wild type yeast (Fig. 4D) (4).

Recent work supports a model in which the Mediator complex, including the kinase module, is first recruited to UAS regions via the tail module, followed by release of the kinase module and association of Mediator with the PIC (4, 18). Our results suggest that TBP plays a critical role in the transit of UAS-associated Mediator to gene promoters. Depletion of TBP together with Kin28 yields a similar Mediator association profile to depletion of TBP alone (Fig. S5), in accord with the notion that TBP is required for a step occurring upstream of Kin28 function, specifically for engagement of Mediator with the PIC. Furthermore, the stronger ChIP signal seen for Med15, from the Mediator tail module, than for Med18, from the head module, at upstream regions, and particularly at SAGA-regulated genes, in *kin28-TBP-AA* yeast is consistent with the tail module being most closely associated with the UAS and the head module being more closely associated with the proximal promoter region (4, 18). Examination of individual gene loci corroborates these observations, with Mediator occupancy shifting upstream at SAGA-regulated genes and often disappearing at TFIID-regulated genes upon TBP depletion (Fig. 5). In some cases Mediator ChIP signal is apparent only after TBP depletion (Fig. 5), indicating that under normal growth conditions transit from the UAS to the promoter occurs too rapidly to allow efficient ChIP of Mediator.

The intermediate effects seen in *kin28-rpb3-AA* yeast may be due to a direct effect on Mediator transit to and association with promoters, or may reflect the impact of loss of Pol II on the cooperative assembly of the PIC. Depletion of Pol II may result in partial loss of TBP, with consequent partial inhibition of Mediator transit from UAS to promoter (44, 49). TBP-containing preinitiation complexes that are present, but which lack Pol II, may facilitate transit of Mediator from the UAS to the promoter, but association may be destabilized by the absence of normal contacts between Mediator and Pol II (27) or other PIC components that are less stably bound in the absence of Pol II.

In sum, our results indicate that PIC components affect the transit of Mediator from UAS to gene promoters, as well as the stable association of Mediator at promoters under conditions that prevent the normal rapid turnover of Mediator that accompanies release of Pol II (i.e., depletion of Kin28).

## Discussion

The work reported here provides novel insight as to how the PIC influences the recruitment and dynamics of Mediator association at Pol II enhancers and promoters. Our major findings are that 1) the Med2-Med3-Med15 tail module triad is dispensable for viability and for Mediator association with active gene promoters; 2) PIC components, in particular Taf1 and Pol II (Rpb3) contribute to Mediator association with active gene promoters, especially at TFIID-regulated genes, and 3) TBP plays a critical role in the transit of Mediator from UAS to promoter sites.

Related to our findings that PIC components facilitate Mediator association with active gene promoters, *in vitro* results have indicated cooperative interactions among Mediator, TFIID, and Pol II (30, 50–52). *In vivo* evidence for such cooperative interactions, however, has been sparse. An “activator bypass” experiment showed that direct recruitment of TFIIB, via tethering to the DNA-binding domain of an activator, resulted in recruitment of a functional PIC and the Mediator complex in yeast, pointing to “backwards” recruitment of Mediator via PIC interactions (53). More recently, depletion of Taf1 was shown to result in a widespread reduction in association of head module subunit Med8 with active genes when mapped by ChEC-seq (8, 54). Neither of these *in vivo* studies, however, examined PIC-Mediator cooperativity at promoter regions, and the distinct effects of depleting different PIC components on Mediator association, with TBP being most important for transit of Mediator from UAS to promoter and Rpb3 and Taf1 being more important for stabilizing occupancy at the promoter following transit (Fig. 4), could not have been predicted on the basis of previous work.

Several interactions between Mediator and TFIID components, and Mediator and Pol II, have been documented that could contribute to Mediator association with promoters. Structural studies have shown the head module subcomplexes Med8-Med18-Med20 and Med11-Med17-Med22 bind with TBP (29, 55). Disruption of the Med8-Med18-Med20 subcomplex in *med18Δ* or *med20Δ* yeast does not cause loss of viability, suggesting that multiple Mediator-TBP interactions are possible at the proximal promoter for initiation (56). In addition, interaction between the tail subunits Med15 or Med16 with TFIID subunits Taf14 or Taf1 have been observed in yeast two-hybrid and mass spectrometry experiments (57–59). Interactions between Mediator and Pol II have also been demonstrated in both structural and genetic studies (3, 24, 27, 55, 60, 61).

Depletion of Rpb3 resulted in effects on Mediator occupancy intermediate between those seen upon depletion of TBP and Taf1 (Figs. 2 & 4). We propose that Pol II contributes both to the stability of the PIC, and specifically to occupancy by TBP, as well as to association of Mediator with the proximal promoter. Thus, depletion of Pol II leads to less efficient transit of Mediator from UAS to the proximal promoter because of partial loss of TBP occupancy, while at those promoters at which Mediator does move from UAS to promoter, Mediator association is rapidly lost because Pol II is not there to stabilize it. In support of a role for Pol II in stabilizing TBP occupancy, Pol II has been previously suggested to function in cooperative assembly of the PIC (30), and depletion of Pol II *in vivo* leads to decreased occupancy of TBP (44, 62).

Based on the results presented here, we propose that Mediator recruitment can occur via two distinct pathways, and that both of these contribute to Mediator function under normal growth conditions (Fig. 6). The first is the canonical recruitment pathway, and is initiated when activator proteins bound to UAS sites engage Mediator through interactions with the Med2-Med3-Med15 triad. UAS-bound Mediator then engages components of the PIC and facilitates stable PIC formation at the proximal promoter (Fig. 6A). This step is accompanied by loss of the CDK module (CKM in the figure) and transit of Mediator from UAS to promoter (4, 18). In the second pathway, PIC components are recruited before or together with Mediator, presumably via activator-mediated interactions, and Mediator association is facilitated by contacts with PIC components and in turn stabilizes binding of those components (Fig. 6B). In both pathways, following formation of the Mediator-PIC complex, TFIIH is recruited and its component subunit Kin28 phosphorylates the carboxyl-terminal domain of the Rpb1 subunit of Pol II; this allows escape of Pol II into the elongation complex, and results in rapid loss of Mediator (20, 21).

**Figure 6.**
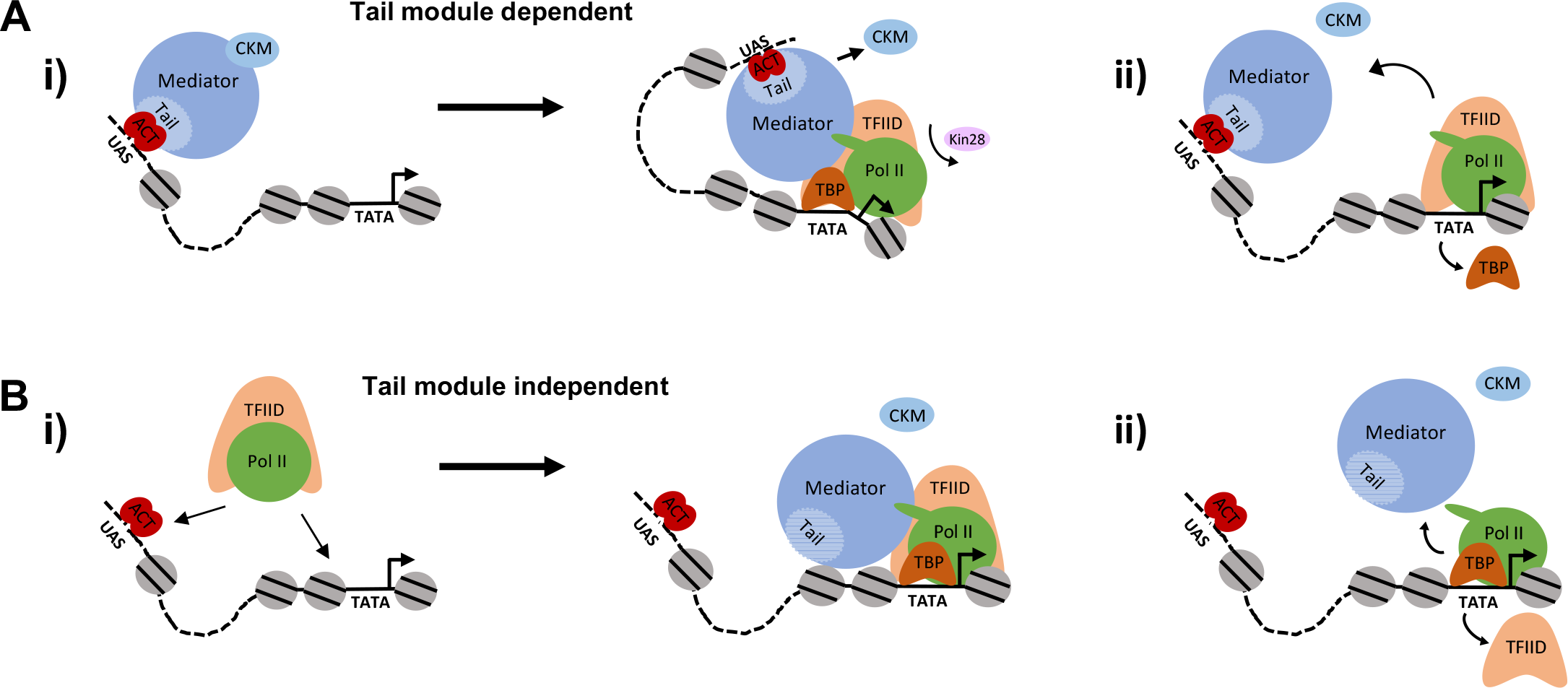
Pathways of Mediator recruitment. (A) (i) Mediator recruitment via interactions between activator proteins bound to UASs and the Med2-Med3-Med15 triad. Contacts between Mediator and components of the general transcription machinery facilitate transit of Mediator to the proximal promoter region, and stabilize association of both Mediator and GTFs. (ii) TBP plays a critical role in this transition, and its depletion results in Mediator being stranded at the UAS. (B) (i) Mediator recruitment via interactions between the middle/head modules and components of the PIC. In this pathway, the PIC is first recruited via interactions between UAS-bound activators and PIC components, such as Tafs within TFIID. (ii) Depletion of PIC components destabilizes association of Mediator with the proximal promoter, particularly at TATA-less, TFIID-regulated promoters. At genes for which interactions between UAS-bound activators and Mediator are insufficiently strong, such depletion results in loss of Mediator association.

We propose that interactions occurring in both pathways contribute to Mediator recruitment at most active genes in yeast. Our results indicate that loss of the tail module triad decreases Mediator association with essentially all active gene promoters (Fig. 1). Furthermore, under appropriate conditions, Mediator association can be detected at more UAS sites than revealed in standard ChIP assays under normal growth conditions. Consistent with this result, Mediator occupancy was reported at UAS regions of essentially all active genes by ChEC-seq (8). Taken together, these findings suggest that Mediator interaction with UAS-bound activators via the Med2-Med3-Med15 triad contributes to Mediator recruitment at the majority of active genes. Similarly, the finding that Mediator association with promoter regions is lost at the majority of genes upon depletion of PIC components (Figs. 4 & 5) suggests that the tail module independent, “PIC-dependent” pathway is also important at many genes. The idea that interactions involving both the tail module and the middle/head modules are important for Mediator association is also consistent with the notion of a single Mediator complex bridging UAS and promoter regions (4, 18).

The tail module independent, PIC-dependent pathway of Mediator recruitment (Fig. 6B) appears more important for TATA-less, TFIID-regulated genes than for TATA-containing SAGA-regulated genes, based on the stronger effect on Mediator occupancy in *med2*Δ *med3*Δ *med15*Δ yeast at the latter class than the former (Fig. 1), and the stronger effect seen at TFIID-regulated genes upon depletion of Taf1 or Rpb3 (Fig. 4). The distinction of these two pathways in terms of dependence on PIC components is also suggested by recent work showing that disruption of a Mediator-TFIIB connection via *med10-ts* yeast differently reduces association of Mediator, TFIIB, and GTFs (28). TFIID-regulated genes exhibit a greater reduction in Mediator rather than GTFs, while SAGA-regulated genes exhibit a greater reduction in GTFs rather than Mediator (28). Furthermore, a recent study reports that Mediator in general, and not just tail module subunits, contributes more strongly to Pol II recruitment at SAGA-regulated than TFIID-regulated genes (63). These findings would be consistent with the first pathway, in which Mediator is recruited via the Med2-Med3-Med15 triad and is critical for subsequent recruitment of GTFs, being most important for SAGA-regulated genes. It should be emphasized, however, that these distinctions reflect general trends, which many individual genes do not follow.

## Experimental procedures

### Yeast Strains

Anchor away (AA) strains used in this study were constructed as described previously and contain the *tor1*-1 mutation and deletion of *FPR1* (38). To allow the AA technique, Taf1, TBP, and Rpb3 were chromosomally tagged with the FRB domain by lithium acetate transformation of PCR products amplified from pFA6a-FRB-kanMX6 into the AA parent strain (yFR1321, generous gift of F. Robert, Institut de Recherches Cliniques de Montréal) (64, 65). Mediator subunits Med2-TAP, Med14-TAP, Med15-myc, Med18-myc, and Med17-myc were tagged by transformation of PCR products amplified from prior strains, as described (17, 44).

To generate double *kin28-taf1-AA* or *kin28-rpb3-AA* strains containing either Med15-myc or Med18-myc, the *kin28-AA* strain (a generous gift of Kevin Struhl, Harvard Medical School) was first switched from *MATα* to *MATa* using the pSC11 plasmid (66), and then mated with the *taf1-AA* and *rpb3-AA* strains, followed by sporulation and isolation of desired products. We were not able to obtain tetrads from *kin28-AA TBP-AA* diploids, and so this strain was constructed by introducing the FRB epitope tag directly into the *kin28-AA* parent strain using standard methods (65). Similarly, *med2*Δ *med3*Δ *med15*Δ yeast were generated by crossing strains LS01H (*med2*Δ) and LS10 (*med3*Δ *med15*Δ) harboring a plasmid bearing the *URA3* gene, followed by sporulation, tetrad dissection, and identification of *med2*Δ *med3*Δ *med15*Δ haploids. A *kin28-AA* strain was constructed from one such segregant by integration of the *tor1-1* mutation, deletion of *FPR1*, and chromosomally tagging *RPL13A* with 2XFKBP and *KIN28* with FRB (38).

Strain genotypes are provided in Table S1, and oligonucleotides used in strain construction are listed in Table S2. For spot dilution assays, yeast cells were grown to an OD_600_ 1.0 and serial dilutions were spotted on YPD medium and incubated at 30°C for 2-3 days, as indicated in the figure legends.

### RNA-seq

RNA was prepared from exponentially growing yeast cultures by the hot phenol method (67). PolyA+ RNA was prepared using a Magnetic Bead Isolation Kit (New England Biolabs), yielding from 1.3% to 9% recovery. Library preparation was performed using the NEB Next Ultra RNA Library Prep kit. Sequencing was performed on the Illumina NextSeq platform at the Wadsworth Center, New York State Department of Health.

### ChIP and ChIP-seq

Conventional ChIP was carried out as described previously (17). Whole cell extracts (WCE) of AA strains were prepared from 50 mL of culture grown in yeast peptone dextrose (YPD) at 30°C to doubling phase at an OD_600_ of 0.8. Rapamycin (LC Laboratories, Woburn, MA) was then added to a final concentration of 1 μg/mL from a 1 mg/mL stock, stored in ethanol at −20°C for not more than one month, for one hour prior to crosslinking. (Concentration of rapamycin stock solutions was determined using A_267_ = 42 and A_277_ = 54 for a 1 mg/ml solution.) Immunoprecipitations were performed using the following antibodies: 5.0 μg Pol II unphosphorylated CTD (8WG16, Biolegend, San Diego, CA), 2 μg c-Myc epitope (9E10, Abcam), 2.5 μg protein A (Sigma), 5 μg TBP (58C9, Abcam, or 2 μL serum, generous gift from A. Weil, Vanderbilt University), and 2.0 μL Taf1 (serum, generous gift from J. Reese and Song Tan, Penn State University).

For ChIP followed by high throughput sequencing (ChIP-seq), library preparation for Illumina paired-end sequencing was performed with the NEBNext Ultra II library prepration kit (New England Biolabs) according to manufacturer’s protocol and barcoded using NEXTflex barcodes (BIOO Scientific, Austin, TX). A size selection step was performed on barcoded libraries by isolating fragment sizes between 200-500 bp on a 2% E-Gel EX agarose gel apparatus (ThermoFisher Scientific). Sequencing was performed on the Illumina NextSeq 500 platform at the Wadsworth Center, New York State Department of Health.

### ChIP-Seq and RNA-seq Analysis

Unfiltered sequencing reads were aligned to the *S. cerevisiae* reference genome (Saccer3) using bwa (68). Up to 1 mismatch was allowed for each aligned read. Reads mapping to multiple sites were retained to allow evaluation of associations with non-unique sequences (68) and the duplicate reads were retained. Binding peaks were identified using SICER (69) with the following parameters: effective genome size 0.97 (97% of the yeast genome is mappable,), window size 50 bp, and gap size 50 bp. Calculation of coverage, comparisons between different data sets, and identification of overlapping binding regions were preceded by library size normalization, and were performed with the “chipseq” and “GenomicRanges” packages in BioConductor (70). Control subtraction was carried out in the following way: coverage (exp)/N1 - coverage (control)/N2, in which “exp” is the data set (in.bam format) to be examined, N1 is the library size of the experimental data (“exp”), and N2 is the library size of the control. Occupancy profiles were generated using the Integrative Genomics Viewer (71). RNA-seq reads were aligned to SacCer3 using tophat2 (72). Non-mRNA species were removed after mapping, and the total number of reads mapping to mRNA species were then calculated and mRNA coverage depth re-normalized. For metagene analysis, including heat maps, reads were normalized against input from *kin28*-AA yeast (KHW127) unless otherwise noted.

RNA-seq and ChIP-seq reads have been deposited in the NCBI Short Read Archive under accession number SAMN07736135-07736171.

## Acknowledgements

This work was supported by grant MCB1516839 to R.H.M, and was also funded in part by the NIH Intramural Research Program at the National Library of Medicine (ZIZ and DL). We are grateful to Todd Benziger for help with strain construction. We thank Kevin Struhl, and Francois Robert for yeast strains; Todd Gray, Tony Weil and Joe Reese for generous provision of antibodies; and Joe Wade for helpful discussions. We gratefully acknowledge help from the Wadsworth Center Applied Genomics Technology and Tissue Culture and Media Cores.

